# Optimization of a Modular Steroid-Inducible Gene Expression System for Use in Rice

**DOI:** 10.1101/595579

**Authors:** Daniela Vlad, Marketa Samalova, Basel Abu-Jamous, Peng Wang, Ian R. Moore, Jane A. Langdale

## Abstract

Chemically inducible systems that provide both spatial and temporal control of gene expression are essential tools, with many applications in plant biology. Using Golden Gate modular cloning, we have created a monocot-optimized dexamethasone (DEX)-inducible *pOp6*/LhGR system and tested its efficacy in rice using the reporter enzyme β-glucuronidase (GUS). The system is tightly regulated and highly sensitive to DEX application, with six hours of induction sufficient to induce high levels of GUS activity in transgenic callus. In seedlings, GUS activity was detectable in the root after *in vitro* application of just 0.01μM DEX. However, transgenic plants manifested severe developmental perturbations when grown on higher concentrations of DEX. The direct cause of these growth defects is not known, but the rice genome contains sequences with high similarity to the LhGR target sequence *lacO*, suggesting non-specific activation of endogenous genes by DEX induction. These off-target effects can be minimized by quenching with isopropyl β-D-1-thiogalactopyranoside (IPTG). The system is thus suitable for general use in rice, when the method of DEX application and relevant controls are tailored appropriately for each specific application.

## INTRODUCTION

The study of gene function in plants relies on reverse genetic tools that facilitate over expression of transgenes or suppression of endogenous gene expression. For some genes, neither approach is feasible due to the development of non-viable phenotypes. To enable functional analysis in such cases, several chemically inducible systems have been established to control transgene expression (for reviews see Zuo and Chua, 2000; Padidam, 2003; Moore at al., 2006; Borghi, 2010). Chemically inducible systems generally consist of chimeric transcription factors and cognate promoters that are derived from genetic elements of heterologous organisms (to avoid interference with expression of endogenous genes) (Moore et al., 2006). The dexamethasone (DEX)-inducible *pOp6*/LhGR gene expression system (Craft et al., 2005; Samalova et al., 2005) is based on a modified *Escherichia coli* lac-repressor system (Moore et al., 1998). The *pOp6* promoter contains a concatemerized binding site comprised of six direct repeats of the 18 base pair *lac* operator (*lacO*) sequence (Craft et al, 2005).This site is bound by the chimeric transcription factor LhGR, which is a fusion between the high affinity DNA-binding domain of the mutant *lac* repressor, *lacI*^His17^, the Gal4 transcription-activation-domain-II and the DEX-binding domain of the rat glucocorticoid receptor (GR) fused at the N-terminus. Addition of DEX to transgenic plants containing both LhGR and *pOp6* leads to relocation of LhGR from the cytoplasm to the nucleus, and consequent transcriptional activation of transgene sequences linked to *pOp6*.

Chemically inducible gene expression systems have been predominantly tested in dicotyledonous species (Zuo and Chua, 2000; Padidam, 2003; Moore at al., 2006; Borghi, 2010). However, two steroid-inducible systems have been tested in rice - the DEX-inducible GVG/*Gal4* (Ouwerkerk et al, 2001) and the estrogen-inducible XVE/*pLex* (Sreekala et al., 2005; Xu et al., 2009; Okuzaki et al., 2011; Chen et al., 2017). Activation of GVG/*Gal4* in rice led to tightly controlled transgene expression but severely perturbed plant growth after exposure to moderate concentrations of inducer. Similar detrimental effects were reported in other species (Kang et al., 1999; Andersen et al., 2003; Amirsadeghi et al., 2007). The XVE/*pLex* system was used more successfully to induce expression in rice callus, roots and leaves but different application methods were required in each case due to inefficient estradiol uptake via the roots. The more recently developed *pOp6*/LhGR system has not yet been tested in monocots but because the *pOp* promoter is not activated by endogenous factors in several species including maize, tobacco and *Arabidopsis* (Baroux et al., 2001; Lexa et al., 2002; Craft et al., 2005; Samalova et al., 2005; Segal et al., 2003, Moore et al., 2006), it may provide a suitable alternative to XVE/*pLex*.

To create a versatile inducible system for transgene expression in monocots, we developed a version of the *pOp6*/LhGR system using the Golden Gate modular cloning system (Weber at al., 2011). The Golden Gate system is designed to provide a rapid, modular and scarless assembly of large constructs, offering a flexible choice of promoters and selection modules. Tests of the system in stable transgenic rice lines revealed tight temporal control over transgene expression and the reliability of a range of approaches for induction.

## RESULTS and DISCUSSION

### Construct design

Three versions of an inducible Golden Gate construct were designed, differing in the choice of selection module and reporter gene. The three versions were designed to test the potential of the *CaMV 35S* promoter (*p35S*) and of any introns present, to influence reporter gene expression. As such, the first construct (17203) contained *p35S* driving expression of the selectable marker gene and a *uidA* derived reporter gene (*GUS*) with two introns, the second (17610) did not contain *p35S* but had introns in the reporter gene, and the third (17613) contained neither *p35S* nor introns in the reporter (Fig. 1). The *pOp6* inducible promoter and the corresponding chimeric transcriptional activator (LhGR), were synthesized as Golden Gate compatible level 0 modules, PU and SC, respectively. *LhGR* was codon optimized for use in rice (*rcoLhGR*) and further ‘domesticated’ to remove all recognition sites for type II restriction enzymes used in Golden Gate cloning: BsaI, BpiI, Esp3I and DraIII (Fig. S1). To test for any unwanted effects caused by codon optimization and/or domestication, the *Arabidopsis* optimised *LhGR2* (Rutherford et al., 2005) and the pOpIn2 bidirectional reporter cassette (Samalova et al., 2019) were cloned into the binary vector pVec8-overexpression (Kim and Dolan, 2016) to obtain construct pVecLhGR (Fig.1, Fig. S2). In all instances the LhGR was expressed under the control of the maize ubiquitin promoter that contains an intron (*pZmUbi*).

**Figure 1.**
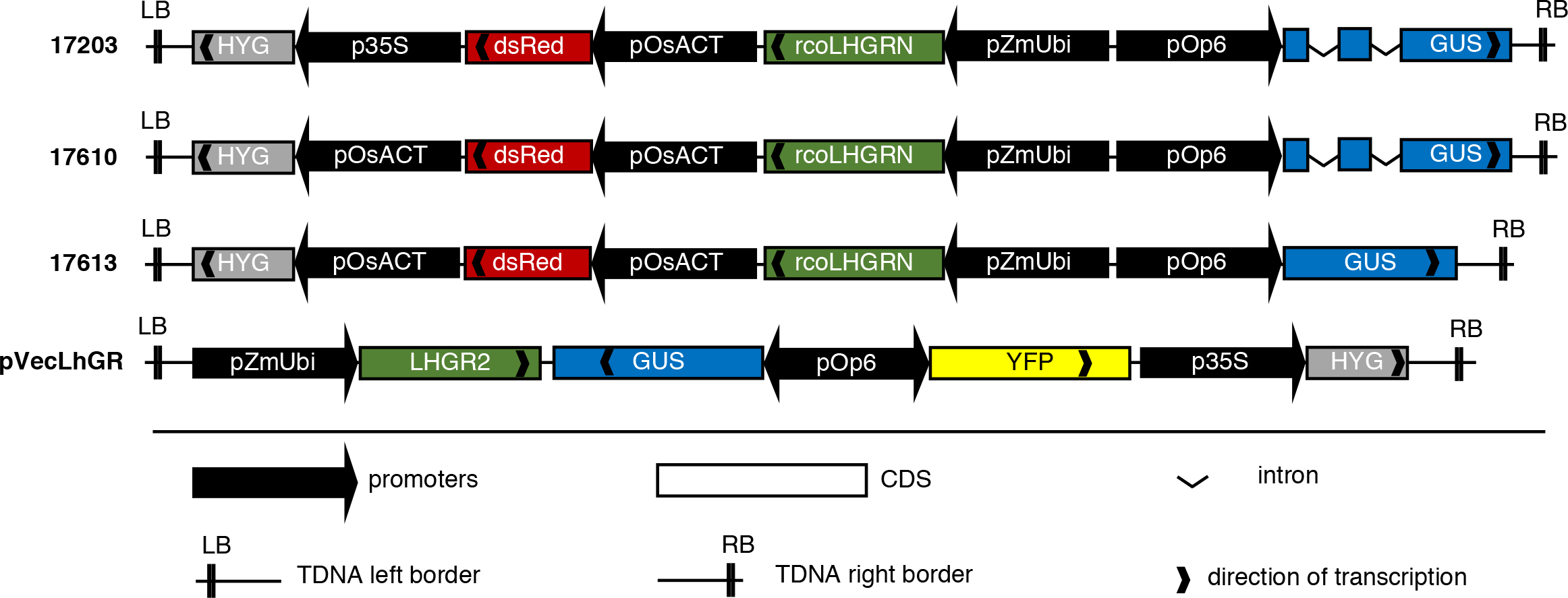
Schematic illustration of the constructs tested in this study. Plasmids 17203, 17610 and 17613 were created using the Golden Gate modular cloning system. Plasmid 17203 contains the selection gene *hygromycin phosphotransferase* (*HYG*) under the control of the *CaMV 35S* promoter (*p35S*), the reporter dsRed gene expressed from the rice actin promoter (*pOsAct*) and a pOp6/LhGR inducible system. The inducible system consists of an activator i.e. a rice codon optimized version (*rco*) of LhGR ubiquitously expressed using the maize ubiquitin promoter (*pZmUbi*), and a *pOp6*:*GUS* reporter. The *GUS* gene contains two introns. Plasmid 17610 differs from 17203, in that the *pOsAct* drives expression of *HYG*. Plasmid 17613 contains an intronless *GUS* gene but is otherwise identical to 17610. All promoter-coding sequence modules contain the *nos* terminator (not represented). The pVecLhGR construct contains the *Arabidopsis* codon optimized *LhGR2* transcriptional activator driven by *pZmUbi* (with the *ocs* terminator), a bidirectional *pOp6* version driving two reporter genes – an intronless *GUS* and yellow fluorescence protein (*YFP*), and *p35S*:*HYG* selectable marker cassette. The construct was assembled as in Figure S2. In all four constructs, the *GUS* reporter gene should only be activated by LhGR upon dexamethasone application.

A control construct was designed to constitutively express an optimized GUS reporter. The *pZmUbi*:*GUSPlus* construct was generated by Gateway cloning of the *GUSPlus* cDNA, amplified from plasmid pCambia1305.2, into the binary destination vector pVec8-Gateway (Kim and Dolan, 2016).

### The pOp6/LhGR system is functional in rice

To test the efficacy of the *pOp6*/LhGR system in monocot species, all constructs were transformed into rice (*Oryza sativa spp japonica* cv Kitaake) to obtain stable transgenic lines. Positive T0 transformants were first validated by histochemical GUS detection after DEX induction of detached leaf fragments, and then transgene copy number was assessed by DNA gel blot analysis (Fig. S3). Lines with one or two copies of the transgene were propagated into the T1 generation. Segregating T1 progeny were tested for GUS activity using an extractable enzymatic assay, after detached leaf fragments were treated with DEX for 24 hours. Notably, most of the positive pVecLhGR T0 lines were silenced, with only ~25% retaining activity in T1 plants, whereas no silencing was observed for any of the other lines.

Compared to activities measured in a constitutive *pZmUbi*:*GUS* line, DEX induction resulted in higher GUS activities in T1 lines transformed with the Golden Gate constructs, but lower activities in the equivalent pVecLhGR transgenic lines (Fig. 2). The rice codon optimized *LhGR* sequence in the Golden Gate constructs might explain the higher rates of induction observed in those lines, but comparisons between lines containing different constructs also suggested that the presence of introns in the reporter gene had an enhancing effect on activity levels (Fig. 2; Fig. S4). There was no correlation between transgene copy number and levels of GUS activity after induction - for example, line 17203_10 was segregating for a single T-DNA insertion and line 17203_7 for two insertions, yet levels of GUS activity after DEX induction were similar in both lines (Fig. 2). Variation in activity between individuals within T1 families may be explained by variable zygosity resulting from segregation, e.g. in line 17613-1 – individual A could be homozygous for transgene insertion whereas B and E could be heterozygous, leading to GUS activity levels twice as high in A than in B and E (Fig. 2).

**Figure 2.**
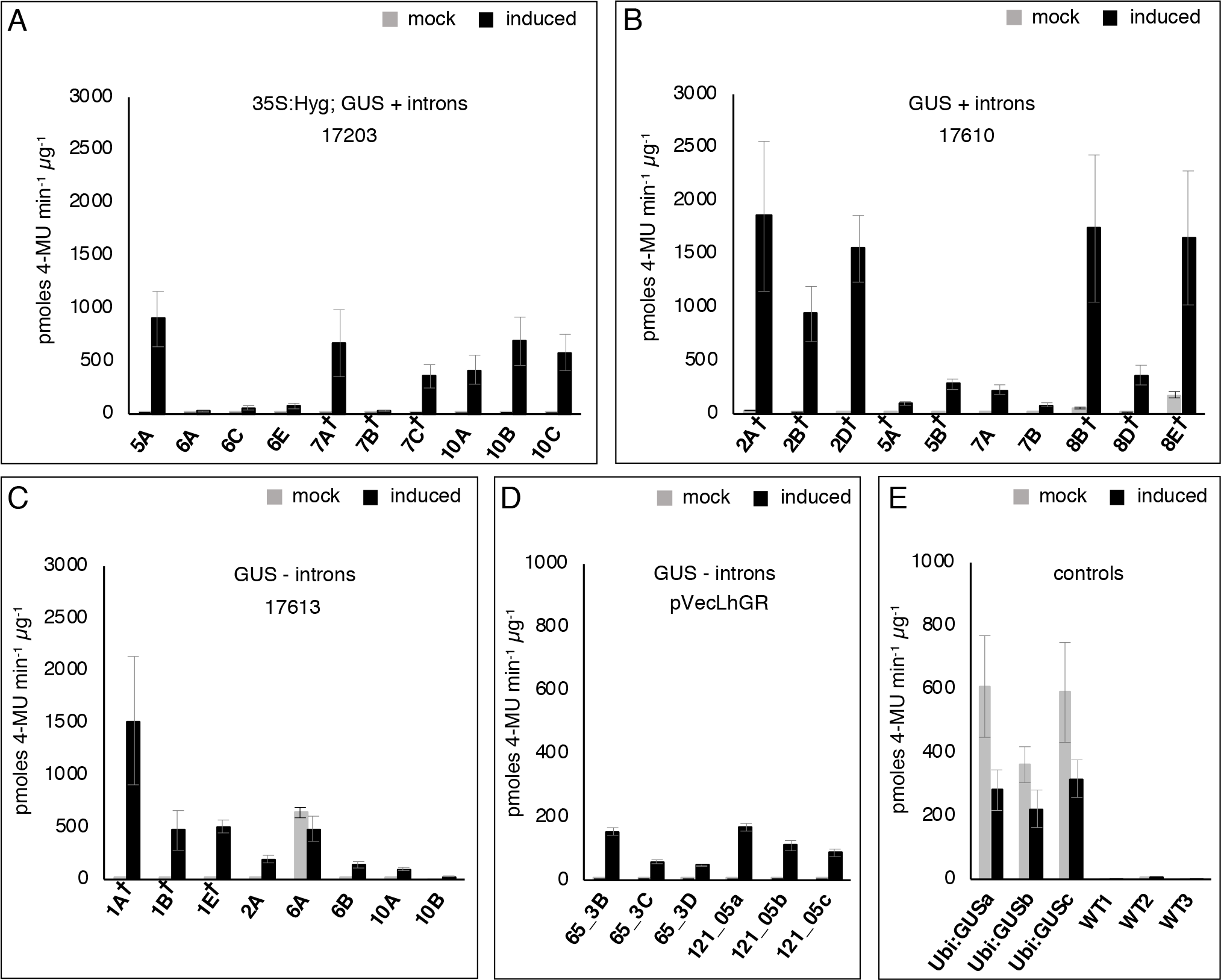
GUS activity is induced in T1 lines. GUS enzymatic activity measured in leaf total protein extracts using 4-methylumbelliferyl ß-D-glucuronide (4-MUG) as a substrate (MUG assay). Bars represent mean ±SD enzymatic activity detected in the absence (grey bars) or presence (black bars) of 10μM DEX. Please note the maximum value on the y-axis is 3000 in plots (A, B, C) and 1000 in plots (D, E). Nomenclature used: number indicates independent transgenic event and letters indicate segregating T1 individuals derived from each event. As such, four independent events were tested for each Golden Gate construct: lines 5, 6, 7 and 10 for 17203, lines 2, 5, 7 and 8 for 17610 and lines 1, 2, 6 and 10 for 17613. † indicates lines segregating for two TDNA insertions, the remaining lines have a single insertion. Two independent pVecLhGR lines (65 & 121) were also tested plus positive *pZmUbi*:*GUS* and negative wild-type (WT) controls.

In some lines, low levels of GUS activity were detected in the absence of the inducer (Fig. 2). In these lines, there was no direct relationship with the number of T-DNA copies integrated into the genome, suggesting that transgene insertion site rather copy number was responsible for “leaky” expression. However, even this explanation is insufficient in the case of line 17613-6, where one T1 individual (A) has very high levels of leaky expression and the other (B) does not. Comparisons between lines 17203/pVecLhGR and 17610/17613 (with and without the *p35S* promoter present in the construct respectively) demonstrate that *p35S* does not drive reporter gene expression in the absence of inducer. However, it remains possible that construct designs that bring the *pOp6* inducible promoter closer to *p35S* might result in transgene activation in the absence of DEX (e.g. as in So et al., 2005).

### Rapid response to dexamethasone application in transgenic rice callus

To test the responsiveness of the *pOp6*/LhGR system to DEX induction, for each construct, callus was generated from a single T2 seed derived from a T1 parental line with only one T-DNA insertion (i.e. 17613_2, 17610_7 and 17203_10). Calli were grown on 10μM DEX for 13 days and sampled at intervals for histological detection of GUS activity (Fig. 3A). No GUS activity was detected in samples prior to DEX treatment. Transgenic callus responded rapidly to DEX exposure, with high levels of GUS activity visible after only 6 hours. Maximum levels of GUS activity were detected after 24 hours in transgenic lines containing constructs with introns in the reporter gene (17203 and 17610) but required 4 days of induction when the intron-less version was present (17613). Activity levels declined between days 10 and 13, even in constitutive *pZmUbi*:*GUS* controls, suggesting that the callus needed sub-culturing at this point to retain viability.

**Figure 3.**
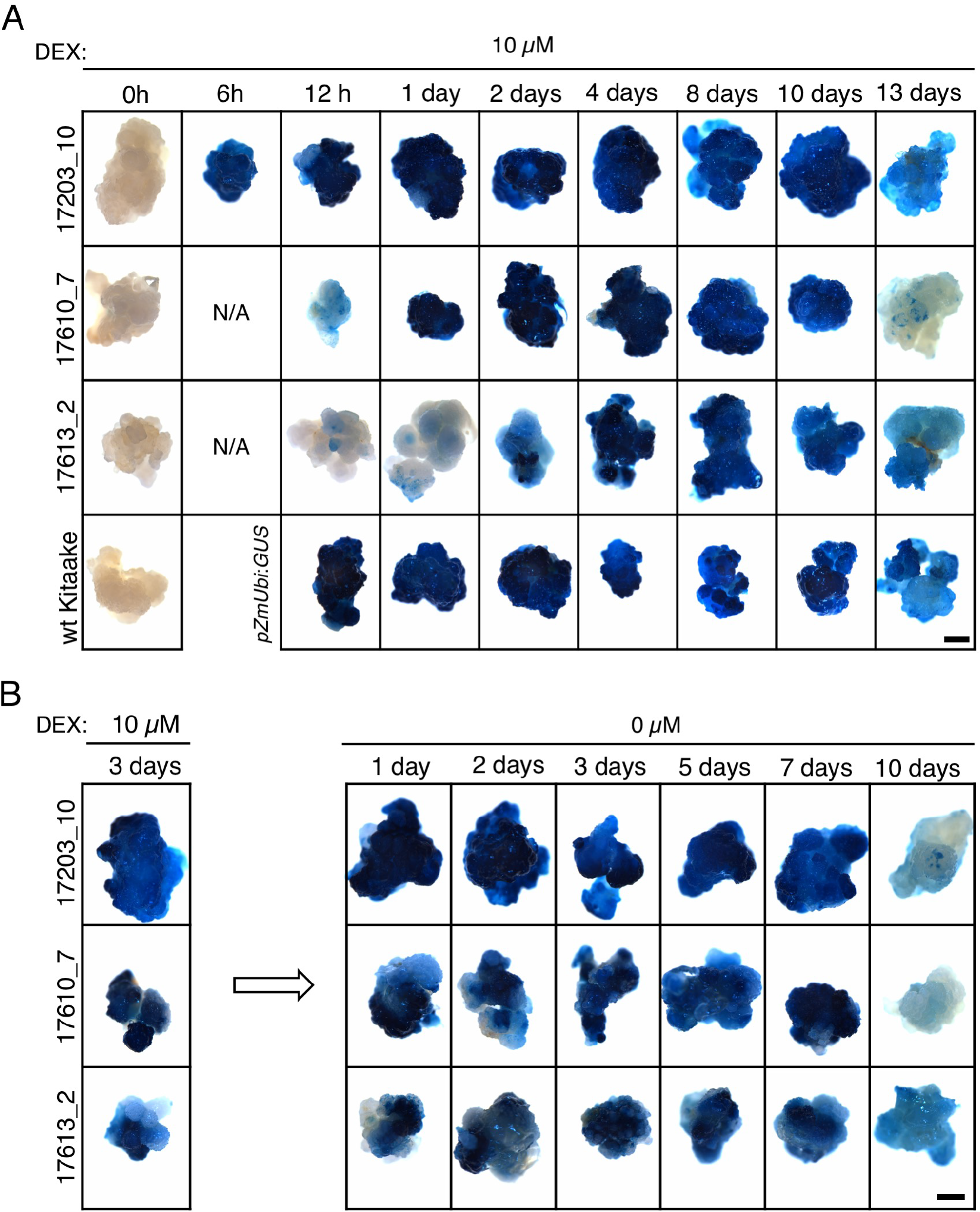
Time course of GUS activity after induction and treatment interruption in transgenic callus. **A, B)** Histological detection of GUS activity in transgenic and wild-type (WT) calli grown for 13 days on solid medium containing 10μM DEX (A) and calli induced with 10μM DEX for 3 days and then transferred to fresh solid medium without DEX for 10 days (B). Scale bars = 1mm.

To determine whether removal of DEX from the culture medium leads to an equally fast response as induction, a reciprocal experiment was performed. Calli induced for 3 days on 10μM DEX-containing media (close to maximum activity level in Fig. 3A) were transferred to fresh ½ MS solid media without DEX and sampled at intervals for a further 10 days (Fig. 3B). GUS expression was maintained at moderately high levels for 7 days and then declined between days 7 & 10. Taking into account the β-glucuronidase expected half-life of ~ 50 hours (Jefferson at al., 1987), GUS activity measurements do not reflect levels of gene expression in real time but rather reflect expression levels approximately 2 days before. As such, high activity levels detected at day 7 reflect expression levels on day 5 after removal from DEX treatment. For faster suppression of gene expression, IPTG could be added to the culture medium at the same time as the inducer is removed (Craft et al.,2005, Schurholz et al., 2018).

### Dexamethasone in the culture medium inhibits growth of transgenic rice seedlings

Developmental studies may require finely tuned manipulation of gene expression at early stages following germination, and in a non-destructive manner that allows phenotypic quantification of transgene effects. To test the effect of applying DEX to seedlings cultured *in vitro*, seeds of *pOp6*/LhGR transgenic lines were germinated and grown on plates containing ½ MS solid media, with and without DEX. All transgenic plants grown on plates without DEX were indistinguishable from wild-type controls (Fig. 4A). However, when exposed to 10μM DEX, with the exception of pVecLhGR lines (Fig. S5), all transgenic lines exhibited a growth arrest phenotype (Fig. 4A). Segregating negative (wild-type) and positive (*pZmUbi*:*GUS*) control lines were not visibly affected by exposure to DEX, indicating that the growth defects were caused by the presence of *pOp6*/LhGR. In this regard, the different behaviour of pVecLhGR lines might be due to less efficient translation and thus lower protein levels of the Arabidopsis optimized *LhGR* versus the rice codon optimized version in the growth inhibited lines.

**Figure 4.**
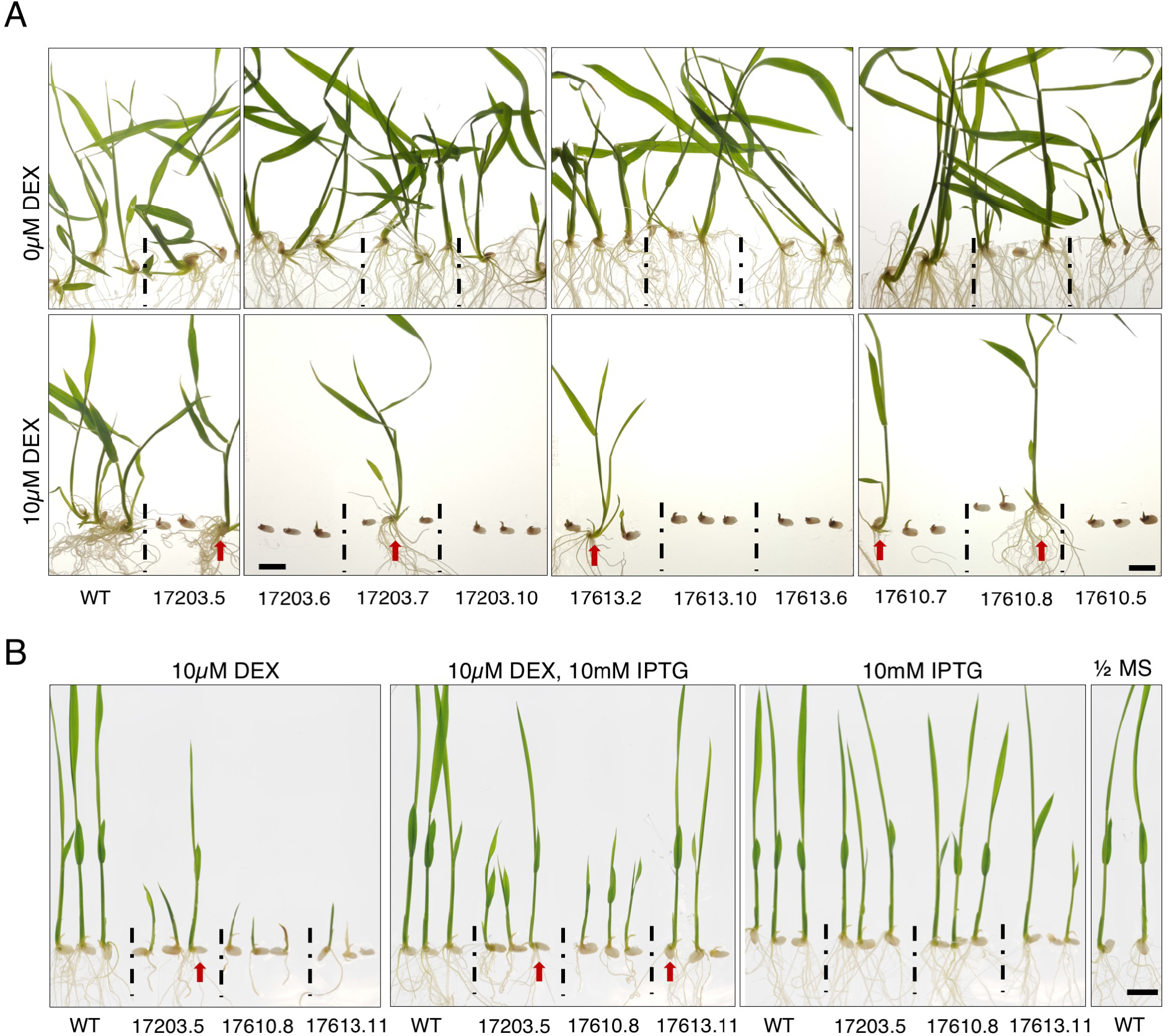
Growth defects caused by exposure to DEX can be ameliorated by addition of IPTG. **A)** T1 plants germinated and grown for 12 days either on ½ MS mock plates containing DMSO or ½ MS plates containing 10μM DEX. **B)** Seeds germinated on ½ MS plates for 3 days were transferred to plates containing either 10μM DEX, 10μM DEX plus 10mM IPTG or 10mM IPTG alone and grown for 5 days. Some transgenic lines are segregating wild-type plants (marked by red arrows) which are not affected by DEX-mediated LhGR activation. Scale bars = 1cm.

Growth defects have not been associated with the use of the *pOp6*/LhGR inducible system in tobacco or Arabidopsis (Samalova et al., 2005; Craft et al., 2005), however, severe growth abnormalities due to likely activation of off-target endogenous genes were reported for the GVG/*Gal4* inducible system in Arabidopsis (Kang et al., 1999), rice (Ouwerkerk et al., 2001), *Lotus japonicus* (Andersen et al., 2003) and tobacco (Amirsadeghi et al., 2007). A similar scenario is plausible for the pOp6/LhGR system in rice. The *lacI* binding domain of LhGR was designed to specifically bind a 18-bp *lacO* sequence in the *pOp6* inducible promoter (Craft et al., 2005). However, *lacI* also has the potential to bind to *lacO* related sequences found in the plant genome (Newell and Gray, 2010). To determine the frequency of potential off target promoters in rice, a bioinformatic analysis was carried out to identify promoters containing *lacO* sequences with 0-5 mismatches. Over 125 potential binding sites were identified (Table1), any of which (depending on the downstream coding sequence) could cause growth defects if binding of LhGR upon DEX induction caused ectopic activation of gene expression. Given that the *lacI* binding site of LhGR can be bound by isopropyl β-D-1-thiogalactopyranoside (IPTG) we tested whether off-target activation could be ameliorated by addition of 10mM IPTG to the DEX-containing growth medium. Figure 4B shows that growth defects were less severe in plants grown in the presence of IPTG, confirming that the perturbations were a direct consequence of LhGR activity, and that off target effects can be quenched.

**Table 1.**
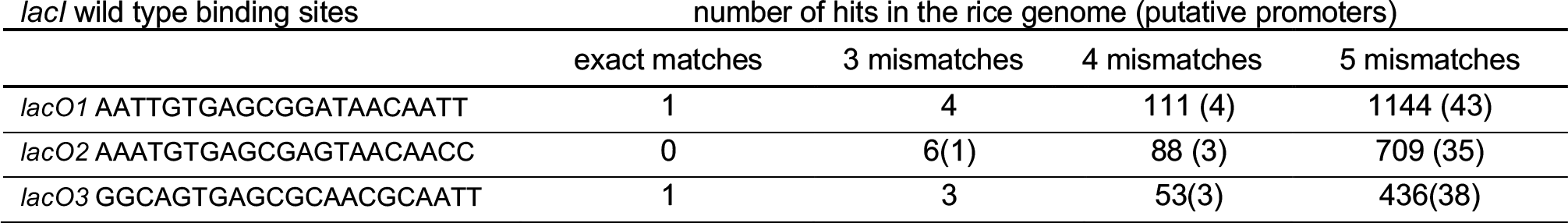
Occurrence of *lacO*-related sequences in the rice genome. The three *E.coli* sequences were used to query the rice genome, with criteria set to return all hits with maximum five mismatches (all of which are predicted to bind *lacI* – based on binding studies in tobacco (Newell and Gray,2010)).

### Transgene expression can be induced by levels of dexamethasone that do not inhibit growth

To try and mitigate the observed inhibitory effect of DEX on plant growth, seeds were pre-germinated on ½ MS plates and then after 3 days transferred to the same medium containing various concentrations of DEX (0.01, 0.1, 1, 5, 10, and 30 μM). Figure 5A shows that the effects of DEX on seedling development were still obvious even in plants grown on concentrations as low as 0.1 μM; the severity of the phenotype, along with levels of detectable GUS activity increasing with the DEX concentration. Importantly, however, low levels of GUS activity could be detected in the absence of any growth inhibition in seedlings transferred to 0.01 μM DEX, suggesting very high sensitivity of the pOp6/LhGR inducible system (Fig. 5A). Because the low levels of transgene expression induced by 0.01μM DEX may not be sufficient for some applications, and higher levels of DEX perturb growth, we tested whether the levels of IPTG used to quench growth defects (Fig. 4B) affected levels of inducible GUS activity. Histological assays carried out on transgenic plants grown in the presence of 10 μM DEX, with or without added 10mM IPTG, revealed similar levels of GUS activity in both conditions (Fig. 5B). This result suggests that for any individual transgenic line, it should be possible to optimize DEX and IPTG concentrations in the culture medium, to minimize growth defects whilst still conferring sufficient levels of induced transgene activity.

**Figure 5.**
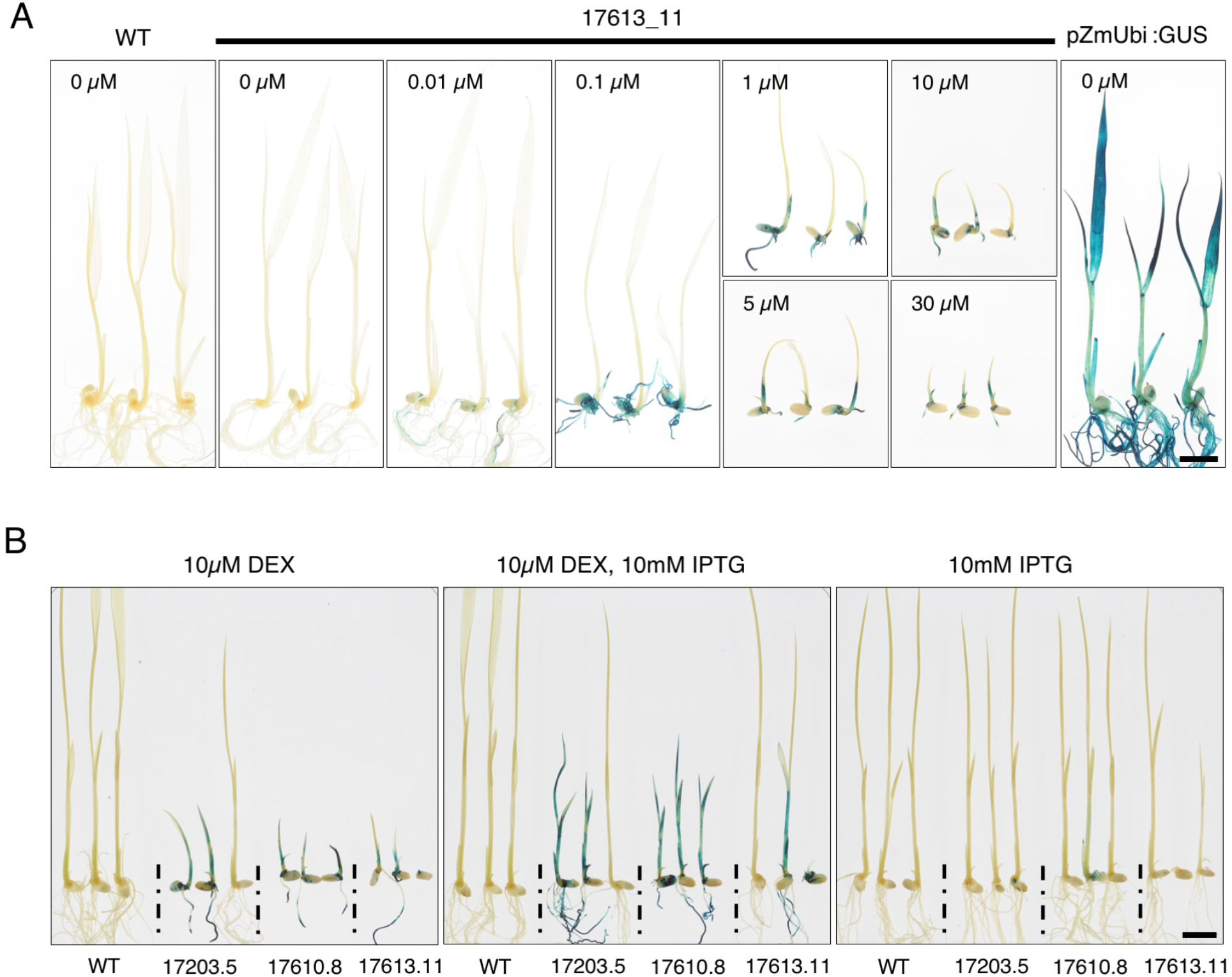
Effects of DEX concentration and IPTG on LhGR mediated activation of gene expression. **A, B)** GUS activity detected in T1 progeny of line 17613_11 grown on plates with 0, 0.01, 0.1, 1,5, 10 and 30μM DEX for 5 days before staining (A) and T1 lines grown on 10μM DEX with and without 10mM IPTG for 5 days (B). Seeds were pre-germinated on ½ MS for 3 days before transferring to treatment conditions. Plants in (B) are the same as those in Figure 4B. Scale bars = 1cm.

To determine whether alternative methods of DEX application can be used to induce transgene expression without inhibiting growth, three further approaches were tested. In the first, 7 day old seedlings that had been grown on ½ MS plates were submerged in a 10 μM DEX solution for ~19 hours before being stained for GUS activity. Figure 6A shows that GUS activity was detected uniformly throughout the transgenic plants, with no apparent morphological differences between the transgenics and wild-type controls. However, transgenic seedlings returned to non-inductive conditions after submergence developed growth abnormalities (Fig. S6). This method is therefore suitable for applications where phenotypes need to be assessed immediately following induction of gene expression.

**Figure 6.**
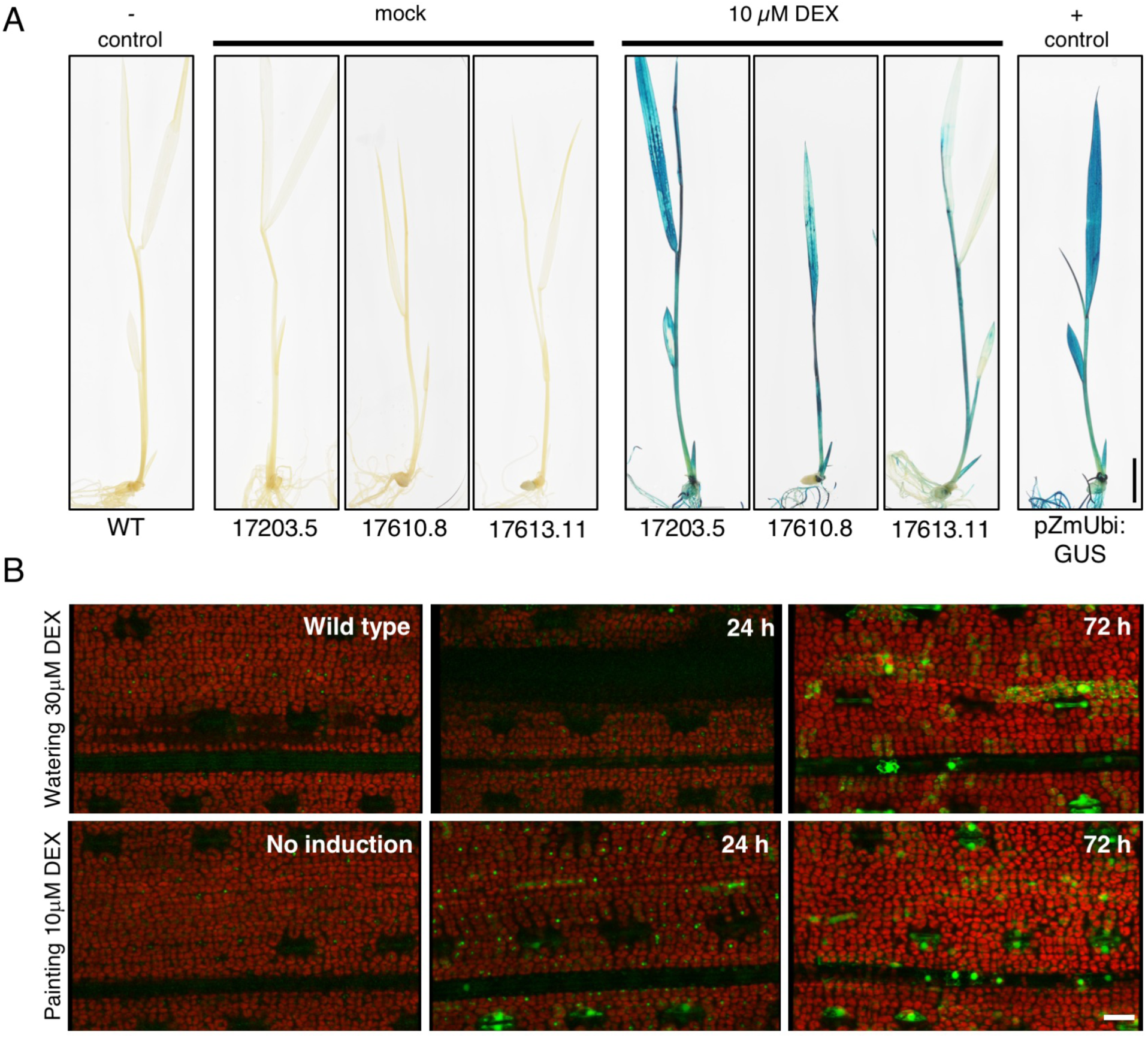
Reporter gene expression induced by seedling submergence, painting or watering. **A)** GUS activity in seven day old seedlings that had been germinated and grown on ½ MS plates before induction by submergence in a 10μM DEX, 0.1% Tween-20 solution for 19 hours. Mock-treated plants (0.1 % DMSO and 0.1% Tween-20) were used as controls. Scale bar = 1cm. **B)** YFP expression imaged under a confocal microscope at 24 and 72 hours following DEX application to leaves of four week old pVecLhGR (line 121C) plants by watering or painting. Scale bar = 20μm.

To test the feasibility of inducing gene expression at later stages of development, either in a systemic or in a localized manner, DEX or the synthetic corticosteroid triamcinolone acetonide (TA) were applied to soil-grown plants by watering or painting. Plants watered with a 30 μM solution at 2 day intervals triggered induction of reporter gene expression at the whole plant level after 72 hours from the time of the first application (Fig. 6B). Local induction of expression was detected after 24 hours when leaves were painted with a 10 μM DEX solution (Fig. 6B) and was restricted to the painted leaf due to minimal transport of DEX between leaves (Craft et al., 2005; Samalova et al., 2005). Higher levels of GUS activity were obtained by watering or painting with TA at equivalent concentrations (Fig. S7). Whole plant and single leaf phenotypes can thus be assessed after induction in this way.

## Conclusion

The optimized *pOp6*/LhGR glucocorticoid-inducible system reported here enables efficient and robust induction of transgene expression in rice. Furthermore, the Golden Gate cloning system used offers the potential to induce multiple transgenes simultaneously with a single LhGR module and to use tissue-specific promoters to spatially control LhGR expression. Induction at low concentrations of DEX and addition of IPTG will suit developmental studies where genes encoding transcription factors do not require high expression levels, and in other applications the use of appropriate controls will serve to distinguish between effects of transgene expression and other changes induced by the presence of LhGR.

## MATERIALS and METHODS

### Generation of recombinant constructs

The *pOp6* inducible promoter and the corresponding chimeric transcriptional activator LhGR-N (referred to as LhGR throughout the manuscript), as reported by Samalova et al. (2005), were adapted for Golden Gate (as below) and synthesized as level 0 modules, PU and SC, respectively. The *LhGR* sequence was codon optimized for use in rice (*rco*) and further ‘domesticated’ to remove all recognition sites for type II restriction enzymes used in Golden Gate cloning: BsaI, BpiI, Esp3I and DraIII.

Cloning was carried out using standard Golden Gate parts and the one-step one-pot protocol (Engler at al., 2008). Golden Gate modules EC47761, EC75111, EC41421, EC47822, EC49283, EC47811 and binary vector pAGM4723 were a gift from Sylvestre Marillonnet and Nicola Patron (ECxxxxx modules are referred to as pICHxxxxx in Weber et al., 2011, Engler et al., 2014); modules EC15069, EC15030, EC15216, EC15455 and EC15073 were a gift from Ben Miller (University of East Anglia, UK) and pICSL4723 was a gift from Mark Youles (The Sainsbury Lab Norwich, UK). All ECxxxxx modules were obtained from the Oldroyd lab (Sainsbury Lab, Cambridge University, UK).

The *pOp6* promoter was cloned into the level 1 backbone EC47761 (position 4, forward) upstream of either a *uidA* (*GUS*) gene containing plant introns (EC75111) or a version with no introns (*kzGUS*, a gift from Dong-Yeon Lee, Donald Danforth Center, St Louis, USA). Modules were finished by addition of a *nos* terminator (EC41421). *LhGR* was cloned downstream of the maize ubiquitin promoter (*pZmUbi*, EC15455) and upstream of the *nos* terminator, into level 1 backbone EC47822 (position 3, reverse).

Level 1 modules described above were assembled into the binary vector pAGM4723 (17203) or pICSL4723 (17610 and 17613) to obtain constructs depicted in Figure 1. Each final construct contained a *hygromycin phosphotransferase* (*HYG*) selection module (EC15069) activated by either a *CaMV 35S* (*p35S*) promoter (level 1 module EC15030) or the rice actin promoter (*pOsACT*, EC15216) in position 1; a *pOsAct*-*dsRed* module (dsRed module EC15073) included to assist with transformed callus selection in position 2 (EC47811); followed by the (*rco*)LhGR module in position3 and a *pOp6* reporter module in position 4. Vectors were closed with end linker ELB4 (EC49283).

To generate the pVecLhGR construct (Fig. 1, Fig. S2), the *LhGR2* and the octopine synthase terminator (*ocs*) sequences were amplified by polymerase chain reaction (PCR) using vector pOpIn2 (Samalova at al., 2019) as the template. The two PCR fragments, *LhGR2* (forward primer P1: 5’-AAAAAGGTACCATGGCTAGTGAAGCTCGAAAAACAAAG-3’; reverse primer P2: 5’-GCATATCTCATTAAAGCAGGACTCTAGTTCACTCCTTCTTAGGGTTAGGTGGAGTATC-3’) and *ocs* (forward primer P3: 5’-TACTCCACCTAACCCTAAGAAGGAGTGAACTAGAGTCCTGCTTTAATGAGATATGC-3’; reverse primer P4: 5’- AAAAAGGTACCCTAAGGCGCGCCGTTGTCGCAAAATTCGCCCTGGACCC -3’) were joined together by overlapping PCR with primers P1 and P4 and cloned into the unique *Kpn*I site of the binary rice overexpression vector pVec8 (Kim and Dolan, 2006). The bidirectional *pOp6* operator array of the pOpIn2YFP vector (generated by M. Kalde, IRM lab) driving expression of GUS and YFP reporters was cloned into a unique *Asc*I site created in the previous cloning step. The correct orientation of the fragment as well as the *LhGR2* in the final pVecLhGR vector was confirmed by sequencing.

The *pZmUbi*:*GUS* construct was generated by Gateway cloning of the *GUSPlus* cDNA into the binary destination vector pVec8-Gateway (Kim and Dolan, 2016). *GUSPlus* cDNA was amplified by PCR from plasmid pCambia1305.2 with Gateway^®^ compatible primers (PW61F: 5’-GGGGACAAGTTTGTACAAAAAAGCAGGCTCATGGCTACTACTAAGCATTTGG-3’; PW61R: 5’-GGGGACCACTTTGT ACAAGAAAGCTGGGTTCACACGTGATGGTGATGG-3’). The coding sequence was subcloned into Gateway^®^ donor vector pDONR™207 in a BP reaction, sequenced, and then cloned downstream of the *ZmUBI* promoter in the binary destination vector pVec8-Gateway (Kim and Dolan, 2016) via an LR reaction.

### Plant transformation

To generate stable transgenic lines, constructs were transformed into *Oryza sativa spp. japonica* cultivar Kitaake using *Agrobacterium tumefaciens*. Calli were induced from mature seeds before co-cultivation with *A. tumefaciens* strain EHA105 that had been transformed with the construct of interest. Callus transformation and seedling regeneration were performed at 32°C according to a protocol modified from Toki et al, (2006) that can be downloaded at https://langdalelab.files.wordpress.com/2015/07/kitaake_transformation_2015.pdf. Regenerated T0 plantlets were verified by polymerase chain reaction (PCR) using primers to detect the presence of the selection gene HYG (forward primer: 5’- CAACCAAGCTCTGATAGAGT-3’; reverse primer: 5’-GAAGAATCTCGTGCTTTCA-3’) and/or by GUS staining, and positive transformants were transferred to soil (John Innes Compost No.2).

### DNA gel blot analysis

Genomic DNA was isolated from 300-400 mg of rice leaf tissue that had been ground in liquid nitrogen using 500μl CTAB extraction buffer (1.5% CTAB, 1.05 M NaCl, 75 mM Tris-HCl, 15 mM EDTA pH8.0) (Murray and Thompson, 1980). After incubating at 65°C for 1 h, samples were thoroughly mixed with equal volumes of chloroform:isoamylalcohol (24:1) and centrifuged at 13000 rpm for 10 min. The aqueous layer was transferred to fresh tubes, precipitated by mixing with equal volumes of isopropanol and centrifuged for 10-15 min at 13000 rpm. Pellets were washed with 70% ethanol, air-dried and dissolved in 50μl dd H_2_O.

For each transgenic plant, 10 μg genomic DNA was digested with the *HindIII* restriction endonuclease. Following electrophoresis in a 1% agarose gel stained with SYBR Safe (Invitrogen), digested DNA was transferred onto Hybond N+ membrane (GE Healthcare, UK). Blots were hybridized with a digoxygenin (DIG)-labelled specific DNA probe for the *HYG* gene and signals were detected using CDP-Star according to the manufacturer’s instructions (Roche Diagnostics)

### Steroid induction of gene expression

Dexamethasone (DEX) was prepared as a 10mM stock solution in dimethyl sulfoxide (DMSO) and stored at −20°C. For each treatment, either DEX (induction) or the equivalent volume of DMSO (mock/control condition) were added to obtain desired concentrations. For application to *in vitro* cultured seedlings, plants were grown on half concentration Murashige and Skoog (1962) medium (½ MS medium) supplemented with 15g/L sucrose. DEX was added to the medium after autoclaving to obtain final working concentrations of 0.01, 0.1, 1, 5, 10 and 30 μM. Before plating, dehulled seeds were surface sterilized with 70% ethanol for 2 min followed by a 15 min wash with a ~2.5% sodium hypochlorite, 0.1% Triton X-100 solution and five washes with sterile water. The plates were incubated with a 16h/8h photoperiod, 30°C day / 25°C night temperatures. Whole seedling induction by submergence was performed in an aqueous solution containing 10 μM DEX and 0.1% Tween-20. Plants grown for 7 days ‘*in vitro’*, as described above, were transferred to 50ml plastic tubes, covered with the induction/mock solution and incubated under the same growth conditions for 19 h.

For callus treatments, callus was obtained from transgenic lines and maintained according to the rice transformation protocol above. To induce GUS activity, DEX was added to the R1 medium at a final concentration of 10 μM.

Isopropyl β-D-1-thiogalactopyranoside (IPTG) was prepared as aqueous 1 M stock solution and added to ½ MS medium after autoclaving to obtain required concentrations. Seeds germinated for 3 days on ½ MS medium were transferred to plates containing 10 μM DEX supplemented with either 1 or 10 mM IPTG.

Gene expression was induced in soil-grown rice by watering and painting with DEX or TA solutions. Four week old rice plants were either watered with 50 ml of 10 μM DEX or newly developed leaves (3^rd^ leaf) were painted on both sides with 30 μM DEX solution supplemented with 0.1% Tween-20. The treatments were repeated at 48h intervals for one week.

### Histological assays of GUS activity

GUS histological detection was performed as previously described (Hay et al, 2006) with minor modifications. Whole seedlings were fixed for 1 h in 90% acetone at −20°C, rinsed in 100mM phosphate buffer, pH 7.6 and stained using 1 mg/ml 5-bromo-4-chloro -3-indolyl-ß-D-glucuronic acid (Melford) supplemented with 0.1% Triton X-100 and 2mM ferrocyanide/ferricyanide salts. Samples were vacuum infiltrated for 10 min before incubation at 37°C for 15 h. Callus samples were fixed for 15- 20 min and then stained for 4 h using the staining solution described above without the addition of Triton-x-100.

### Enzymatic assays of GUS activity (MUG assay)

GUS (*ß- glucuronidase*) enzymatic activity was measured based on a method described by Jefferson et al, (1987). Extracts were prepared from ground leaf samples using 10ul protein extraction buffer per mg fresh tissue. Total protein concentration was determined using the Bio-Rad Protein Assay according to the manufacturers protocol for microtiter plates.

The fluorogenic reaction was carried out in 96 well-plates (FLUOTRACtm 200) at 37°C using protein extraction buffer supplemented with 4-methylumbelliferyl ß-D-glucuronide (4-MUG) as a substrate. The fluorophore 4-methyl umbelliferone (4-MU) produced upon 4-MUG hydrolysis by ß- glucuronidase (GUS) was quantified by measuring emission at 455 nm using a FLUOstar Omega Microplate Reader set at 365 nm excitation wavelength. A standard curve of 4-MU was used to calculate the amount of 4-MU/unit of time. Wild type rice protein was added to the standard curve to correct for eventual autofluorescence or quenching. Activity was calculated from three technical replicates and expressed in (pmoles 4-MU) min^−1^ (μg protein)^−1^.

### Detection of YFP fluorescence

YFP signal was detected using a Zeiss LSM 510 Meta laser-scanning microscope with 514-nm excitation from an Argon laser and BP535-590IR emission filter.

### Bioinformatic screening for the presence of *lacO*-related sequences

The rice reference genome v7.0 was scanned for sequences matching any of the three *lacI* wild type binding sites, with up to 5 mismatches. A brute force scan was adopted with a sliding window equal in length to the searched sequence that slides one base at every step of scanning. At any base of the reference genome, if the reverse complement of the searched sequence matched the forward strand, the sequence genome going backwards from that base was counted as a match on the reverse strand. To find promoter regions, genes in proximity of the detected matches were identified. If an annotated gene was on the same strand as the match and started within 1000 bases downstream of the match’s first base, then the match was defined as being a putative promoter sequence.

## AUTHOR CONTRIBUTIONS

DLV, MS, IRM and JAL conceived and designed the study; DLV carried out the cloning, transformation and analysis of Golden Gate lines; MS created and analysed the pVecLhGR lines; PW cloned the pZmUbi:GUS construct and generated the pZmUbi:GUS transgenic lines; BA-J performed the bioinformatic screen; DLV and JAL wrote the first draft manuscript; all authors had input to the final version.

## ACKNOWLEDGEMENTS

We are grateful to Sylvestre Marillonnet, Nicola Patron, Ben Miller, Mark Youles and Dong-Yeon Lee for providing Golden Gate modules used in this study and John Baker for photography. The work was funded by a C_4_ Rice Project grant from The Bill & Melinda Gates Foundation to the University of Oxford (OPP 1129902).

## SUPPORTING INFORMATION

**Figure S1.** Golden Gate compatible pOp6/LhGR level 0 modules

**Figure S2.** Cloning scheme for the pVecLhGR construct

**Figure S3.** DNA gel blot analysis of transgenic lines.

**Figure S4.** GUS enzymatic activities measured in T0 lines.

**Figure S5.** Phenotype of T1 pVecLhGR line 121.

**Figure S6.** Effect of induction by submergence in transgenic seedlings.

**Figure S7.** GUS enzymatic activities measured in pVec-LhGR lines induced in soil.

**Table S1.** Summary of T-DNA insertion numbers in each transgenic line tested.

