## Supporting Information for "Optimization of a Modular Steroid-Inducible Gene Expression System for Use in Rice"

### SUPPLEMENTARY INFORMATION

**A**

>pL0M-PU-pOp6

***ggag***AATTGTGAGCGCTCACAATTGAAAGACTAGAAAGAAGAAAGGGAAGAGAAAGAATTGTGAGCGCTCA
CAATTGAAAGACTAGAAAGAAGAAAGGGAAGAGAAAGAATTGTGAGCGCTCACAATTGAAAGACTAGAAAG
AAGAAAGGGAAGAGAAAGAATTGTGAGCGCTCACAATTGAAAGACTAGAAAGAAGAAAGGGAAGAGAAAGA
ATTGTGAGCGCTCACAATTGAAAGACTAGAAAGAAGAAAGGGAAGAGAAAGAATTGTGAGCGCTCACAATT
GAAAGACTAGTGGATCGATCTTCGCAAGACCCTTCCTCTATATAAGGAAGTTCATTTCATTTGGAGAGGAC
ACG***aatg***

**B**

>pL0M-SC-rcoLhGR

***aatg***GCATCAGAGGCACGTAAGACTAAGAAGAAGATCAAAGGAATTCAGCAGGCAACAGCAGGAGTGAGCC
AAGACACCTCTGAGAATCCGAATAAGACGATAGTGCCGGCCGCTCTGCCCCAACTGACTCCGACACTTGTG
TCACTGCTGGAGGTAATAGAGCCGGAGGTCTTGTATGCCGGCTACGATTCGTCCGTCCCTGACAGTGCCTG
GCGCATTATGACCACGCTCAACATGCTCGGTGGGCGGCAAGTCATTGCGGCGGTTAAGTGGGCGAAAGCAA
TTCCTGGTTTCAGGAATCTCCATCTCGATGACCAAATGACCCTCCTCCAATACTCCTGGATGTTCCTGATG
GCGTTTGCCCTCGGTTGGCGCAGTTACCGCCAATCAAGCGGAAACTTGCTGTGCTTTGCACCTGACTTGAT
CATCAATGAACAGAGGATGTCGCTGCCGTGCATGTACGACCAGTGCAAACATATGCTCTTCGTGTCCTCCG
AATTGCAGCGCCTTCAAGTGAGCTACGAAGAATACTTGTGCATGAAAACTCTGCTCTTGTTGAGTTCCGTG
CCTAAGGAGGGACTCAAGAGCCAAGAGCTGTTTGATGAAATCAGGATGACATACATCAAGGAGCTCGGGAA
AGCGATCGTAAAGCGCGAGGGTAACTCATCACAGAACTGGCAACGCTTCTACCAGCTGACTAAGTTGCTCG
ATTCCATGCATGAGGTAGTCGAAAACCTCCTGACCTATTGCTTCCAAACCTTCCTCGATAAGACCATGTCT
ATTGAGTTTCCCGAGATGCTCGCAGAGATAATCACGAACCAGATCCCCAAGTACAGCAACGGTAATATAAA
GAAGCTTCTCTTCCATCAGAAATCTACCTCTAAACCAGTGACCTTGTATGACGTGGCGGAGTACGCCGGAG
TCAGTCATCAAACAGTTTCAAGGGTGGTTAACCAGGCGTCCCACGTTTCGGCAAAGACCAGAGAAAAAGTG
GAGGCAGCTATGGCGGAACTGAATTACATTCCAAACCGCGTGGCACAACAACTGGCGGGAAAGCAAAGCCT
CCTTATTGGGGTGGCAAC***GTCTTC***CCTCGCTCTGCATGCCCCTTCTCAGATTGTTGCTGCCATAAAGAGTC
TCC *(Ser)*
GCGCAGACCAGCTCGGTGCAAGTGTGGTTGTTTCCATGGTCGAAAGGTCGGGTGTGGAGGCGTGTAAAGCG
GCCGTGCACAATTTGCTCGCCCAACGCGTTTCGGGCCTCATAATTAATTACCCTCTGGACGATCAGGATGC
GATTGCCGTAGAAGCAGCTTGTACGAACGTCCCGGCGCTCTTCTTGGACGTCAGCGACCAAACCCCGATTA
ATTCAATTATATTCAGCCATGAGGACGGGACG***CGTCTC***GGCGTGGAGCACCTCGTAGCGCTGGGACATCAA
CGC *(Arg)*
CAAATAGCGCTTCTGGCTGGTCCGCTCTCTTCTGTCTCGGCTCGGCTTAGGCTTGCTGGGTGGCATAAGTA
TTTGACACGTAATCAGATCCAACCAATCGCGGAGAGGGAGGGCGACTGGAGTGCTATGTCTGGATTCCAAC
AGACTATGCAAATGCTCAATGAGGGAATAGTGCCTACCGCCATGCTGGTGGCCAATGATCAAATGGCACTT
GGAGCGATGCGCGCGATTACCGAATCAGGGCTTAGAGTCGGGGCTGACATCTCTGTCGTGGGGTATGATGA
CACTGAGGACTCCTCGTGTTATATCCCACCTCTTACGACAATAAAACAGGACTTC***CGTCTTC***TGGGGCAGA
CGC *(Arg)*
CGTCAGTCGACCGCCTTCTTCAACTGAGCCAGGGTCAGGCGGTAAAGGGCAACCAACTCCTGCCCGTGTCT
CTGGTGAAGCGTAAGACTACAAGTGGCTCAGAATTCGCTAATTTCAATCAGTCCGGGAATATAGCGGACTC
GAGCCTTTCCTTCACTTTTACGAACTCGAGTAACGGCCCTAACCTCATCACCACACAAACAAACAGCCAGG
CCCTCTCGCAGCCGATTGC***GTCTTC***AAACGTCCACGATAATTTCATGAACAATGAAATTACCGCTAGCAAA
TCC *(Ser)*
ATAGATGATGGGAACAATTCAAAACCACTGTCACCAGGGTGGACAGATCAGACCGCTTACAACGCTTTCGG
TATCACGACGGGCATGTTTAATACCACTACCATGGACGACGTCTACAATTACCTGTTTGACGATGAGGACA
CACCACCTAATCCGAAGAAGGAGTGA***gctt***

**Figure S1. Golden Gate compatible pOp6/LhGR level 0 modules A)** The pOp6 inducible promoter contains six *lac operator* sequences highlighted in grey. Sequence is flanked by the level 0 PU fusion sites, ***ggag/aatg***. **B)** Rice codon optimized version of the chimeric transcription activator LhGR with recognition sites for *BpiI* and *Esp3I* marked in bold italics. Codons are underlined and the domesticated versions containing in each case a silent base pair change are marked in red. The amino acid encoded by each triplet is specified in brackets. Sequence is flanked by the level 0 SC fusion sites, ***aatg/gctt.***

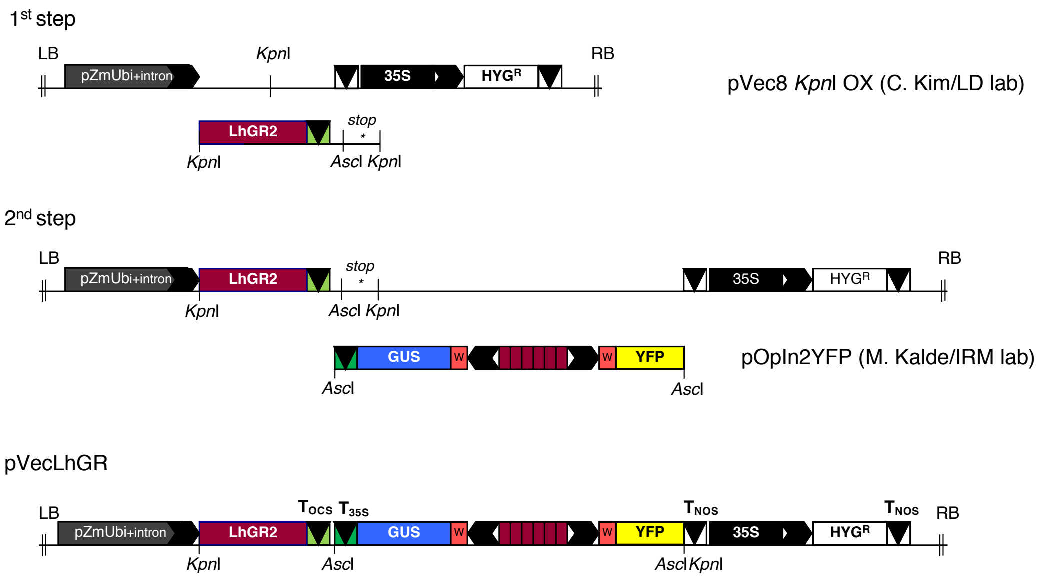


**Figure S2. Cloning scheme for the pVecLhGR construct**

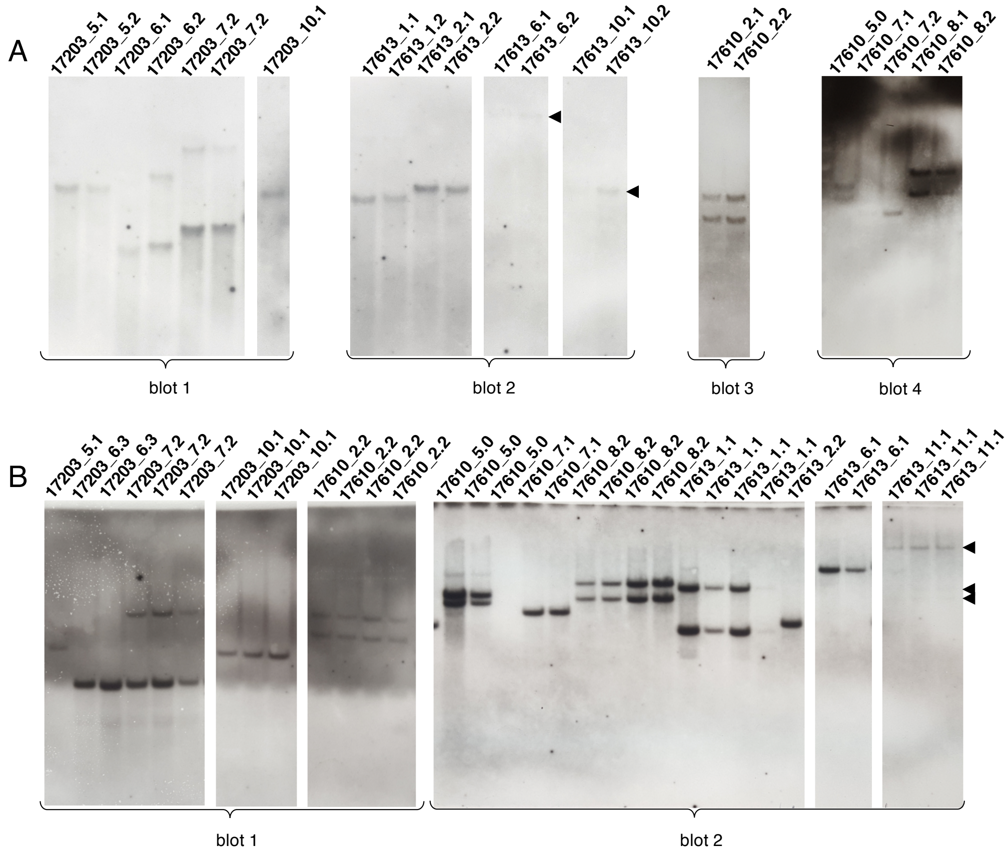


**Figure S3. DNA gel blot analysis of transgenic lines. A, B)** Number of T-DNA inserts detected following genomic DNA digestion with *HindIII* and hybridization with a DIG-labelled HYG probe in T0 transgenic plants (A) and their progeny (B). In (A), numbers identify T0 plants resulting from the same transformation event (e.g. 17203_5.1 and 5.2) and in (B) segregating individuals labeled with same the parental line number (e.g. 17203_7.2) are segregating progeny form that line

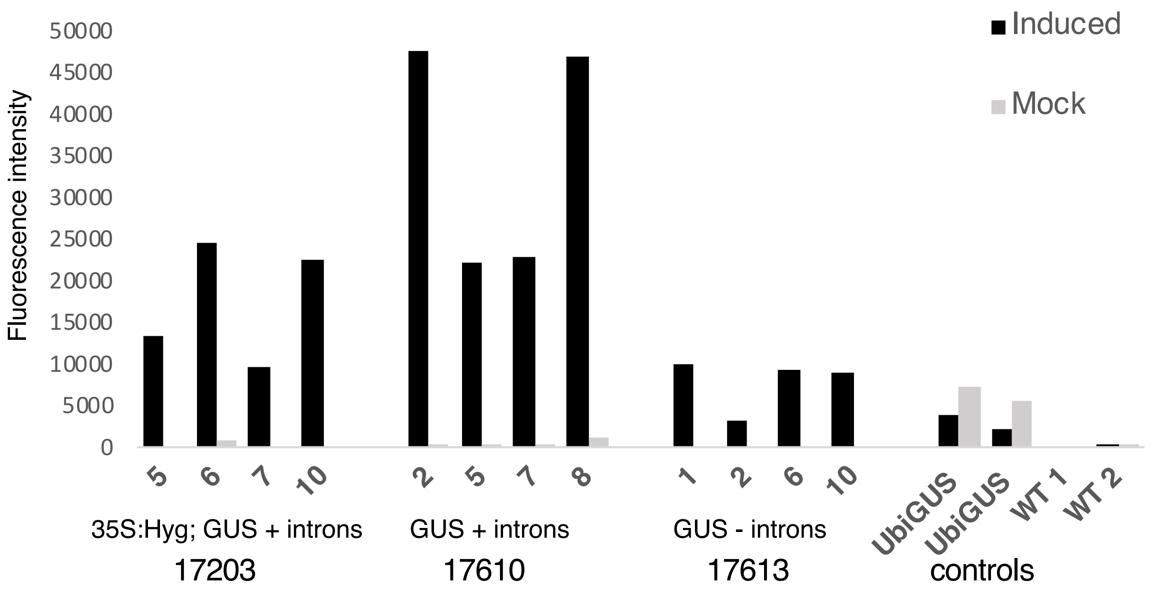


**Figure S4. GUS enzymatic activities measured in T0 lines.** Enzymatic activity was first tested in a MUG assay using 50µl protein extracted from equal amounts of leaf tissue. Bars represent fluorescence intensities measured for non-induced (grey bars) or DEX-induced (black bars) samples.

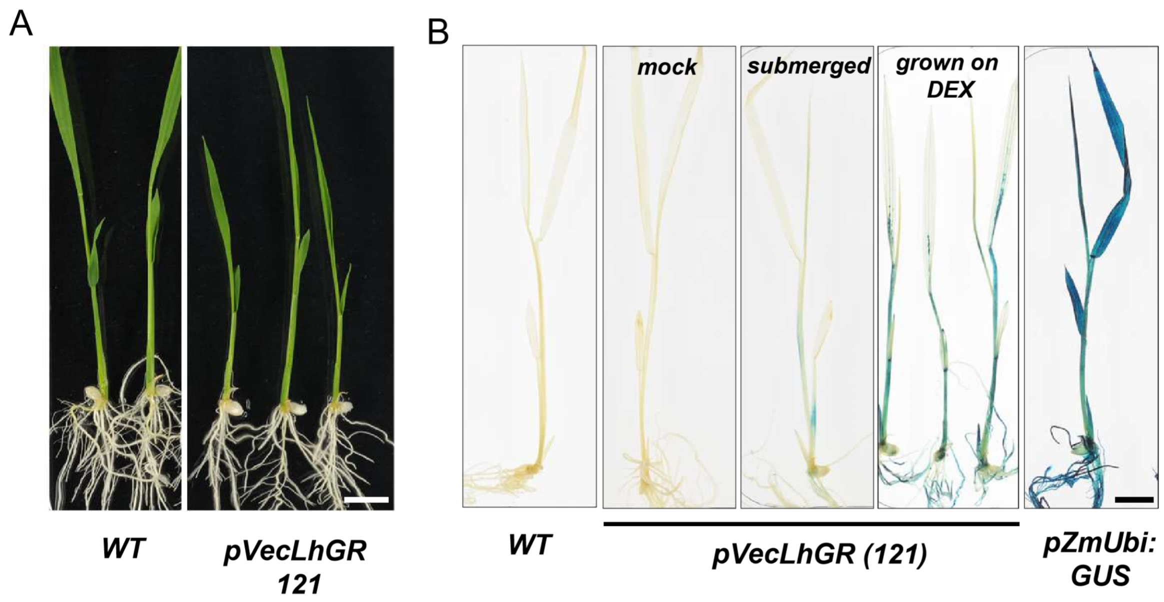


**Figure S5. Phenotype of T1 pVecLhGR line 121. A)** T1 progeny of line 121 germinated on ½ MS for 3 days and grown on 10µM DEX for 5 days. **B)** GUS activity in T1 seedlings derived from line 121 induced in the presence of 10µM DEX applied either by submergence or through the '*in vitro'* growth medium. Scale bars = 1 cm

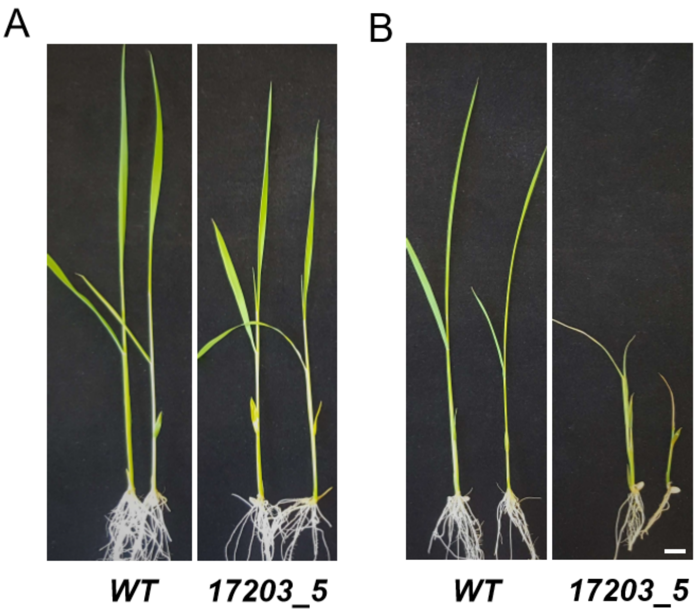


**Figure S6. Effect of induction by submergence in transgenic seedlings. A, B)** Phenotype of seedlings that were induced by submergence in a 10μM DEX solution and then transferred to hydroponic growth conditions for 5 days. Control (not submerged) seedlings develop normally (A) whereas abnormal growth is observed in transgenic seedlings (B). Scale bar = 1 cm.

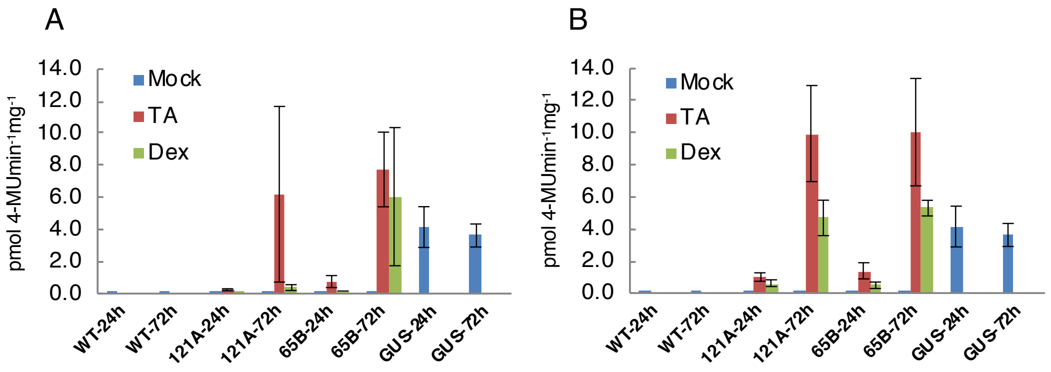


**Figure S7. GUS enzymatic activities measured in pVec-LhGR lines induced in soil.** Four week old rice plants were induced either with DEX or triamcinolone acetonide (TA) by watering (A) or painting (B) with 30μM or 10μM solutions respectively. Leaves were sampled and enzymatic activity measured in a MUG assay 24 and 72 hours following first application of the inducer.

| lines | TDNA insertions in T0 | TDNA insertions in T1 |
|---|---|---|
| 17203_5.1 | 1 | 1 |
| 17203_6.3 | 1 | 1 |
| 17203_7.2 | 2 | 2 (linked) |
| 17203_10.1 | 1 | 1 |
| 17610_2.2 | 2 | 2 (linked) |
| 17610_5.0 | 2 | 2 (linked) |
| 17610_7.1 | 1 | 1 |
| 17610_8.2 | 2 | 2 (linked) |
| 17613_1.1 | 2 | 2 (linked) |
| 17613_2.2 | 1 | 1 |
| 17613_6.1 | 1 | 1 |
| 17613_10.2 | 1 | 1 |
| 17613_11.1 | N/A | 3 |

**Table S1. Summary of T-DNA insertion numbers in each transgenic line tested.** Constitutive pZmUbi:GUS lines were used as controls. Three plants from segregating populations were used for each treatment. The error bars represent SD.
